# Heterologous Expression of NoxA Confers Aerotolerance in *Clostridium sporogenes*

**DOI:** 10.1101/2022.05.20.491843

**Authors:** Sara Sadr, Bahram Zargar, Justo Perez, Marc Aucoin, Brian Ingalls

**Affiliations:** Department of Chemical Engineering, University of Waterloo, Waterloo, ON, Canada; Department of Applied Mathematics, University of Waterloo, Waterloo, ON, Canada; CREM Co, 3403 American Drive, Mississauga, ON, Canada

**Author notes:** **Correspondence:** Brian Ingalls.

**Keywords:** *Clostridia*, *C. sporogenes*, aerotolerance, NADH oxidase, *noxA* gene

## Abstract

*Clostridium* is a genus of Gram-positive obligate anaerobic bacteria. Some species of *Clostridium*, including *C. sporogenes*, may be of use in bacteria-mediated cancer therapy. Spores of *Clostridium* are inert in healthy normoxic tissue, but germinate when in the hypoxic regions of solid tumors, causing tumor regression. However, such treatments fail to completely eradicate tumors because of higher oxygen levels rise at the tumor’s outer rim. In this study, we demonstrate that a degree of aerotolerance can be introduced to *C. sporogenes* by transfer of the *noxA* gene from *C. aminovalericum*. NoxA is a water-forming NADH oxidase enzyme. Thus, its activity has no detrimental effect on cell viability. In addition to its potential in cancer treatment, the *noxA*-expressing strain described here could be used to alleviate challenges related to oxygen sensitivity of *C. sporogenes* in biomanufacturing.

## 1 Introduction

*Clostridium* is a genus of Gram-positive obligate anaerobic bacteria. The *Clostridium* species *C. sporogenes* has attracted attention for potential use in treating solid-mass tumors (Fox *et al*., 1996; Kubiak and Minton, 2015). Spores of *Clostridium* are inert in normoxic healthy tissue, but germinate when in the hypoxic regions of solid tumors (Mowday *et al*., 2016). It has been shown that *C. sporogenes* exhibits accurate targeting of necrotic cores (Lemmon *et al*., 1997; Kubiak and Minton, 2015). However, the oxygen sensitivity of *C. sporogenes* ultimately limits its applicability to tumor treatment. *C. sporogenes* fails to completely eradicate tumors because it cannot penetrate the outer edge of tumors where the oxygen level is higher (Mengesha *et al*., 2007; Mowday *et al*., 2016).

Despite being characterized as obligate anaerobes, sensitivity to oxygen varies widely within the *Clostridium* genus, with individual species demonstrating an array of different mechanisms for dealing with the presence of oxygen. Some species of *Clostridia* undergo endosporulation to resist oxygen stress (Diallo, Kengen and López-Contreras, 2021; Morvan *et al*., 2021). In contrast, *C. butyricum* expresses NADH and NADPH oxidase in the presence of oxygen (Kawasaki *et al*., 1998). *C. aminovalericum* is equipped with an oxidase gene, *noxA*, which is strongly expressed when the bacteria are exposed to oxygen (Kawasaki *et al*., 2005). In *C. acetobutylicum*, many oxygen responsive genes have been identified, such as the *nror* gene cluster, which enables oxidoreductase activity (Kawasaki *et al*., 2005, 2009). Of the *Clostridium* species, *C. sporogenes* has been reported to be one of the least aerotolerant (Kawasaki *et al*., 1998).

We hypothesized that a degree of aerotolerance could be introduced to *C. sporogenes* by transfer of the *noxA* gene from *C. aminovalericum*. NoxA is a water-forming NADH oxidase, therefore its activity should have no detrimental effect on viability. As reported below, we found that heterologous expression of *noxA* gene allows *C. sporogenes* to better tolerate oxygen stress. We propose that this engineered strain, which we refer to as PTN, could be used in tumor treatment; it presents a mechanism for tuning aerotolerance to allow growth in the outer rim of a solid tumor.

In addition to its potential applications in cancer therapy, *C. sporogenes* has also been investigated as a means of biofuel production (Gottumukkala *et al*., 2013, 2015). In that context, oxygen sensitivity presents challenges in maintaining anaerobic culture environments, resulting in unsatisfactory cell densities (Papoutsakis, 2008; Zeng, 2019; Zhao *et al*., 2020). The *noxA*-expressing strain described here could be useful in that context.

## 2 Materials and Methods

### 2.1 Bacteria Strains and Plasmids

*Clostridium sporogenes* NCIMB 10696 (referred to herein as the native strain), and *Escherichia coli CA434* were gifts from Professor Nigel Minton (University of Nottingham, UK) and Professor Mike Young (Aberystwyth University, UK), respectively. The shuttle vector pMTL825x was a gift from Prof. Minton. The genome of *C. aminovalericum* (ATCC 13725) was purchased from the American Type Culture Collection (<u>VA, USA</u>). All restriction enzymes were purchased from New England Biolabs (Whitby, ON, Canada). The plasmid pGlow-Xn-Pp1-CI was purchased from BioCat GmbH (Heidelberg, Germany).

To construct the transformed *C. sporogenes* strain, which we refer to as PTN, the *thl* promoter was PCR-amplified from the pGlow plasmid with primers thl-f and thl-r (Table 1). The PCR product was cloned into a T Easy plasmid using the pGEM®-T Easy Vector (Promega, USA), following the manufacturer’s protocol. The T Easy plasmid was digested using EcoR1 and SpeI and the resulting insert was cloned into the EcoR1-XbaI site of the pMTL825x shuttle vector to construct pMTL_thl. The *noxA* gene from *C. aminovalericum* (GenBank sequence AB219226.1) was PCR-amplified with primers noxA-XhoI-F and noxA-NheI-R (Table 1). The PCR product was cloned into a CloneJET vector (CloneJET PCR Cloning Kit, Thermo Scientific, USA) following the manufacturer’s protocol. The clonejet_noxA plasmid was digested by NheI and XhoI and the resulting insert was cloned into the XhoI-NheI sites of the pMTL_thl plasmid.

**Table 1-.**
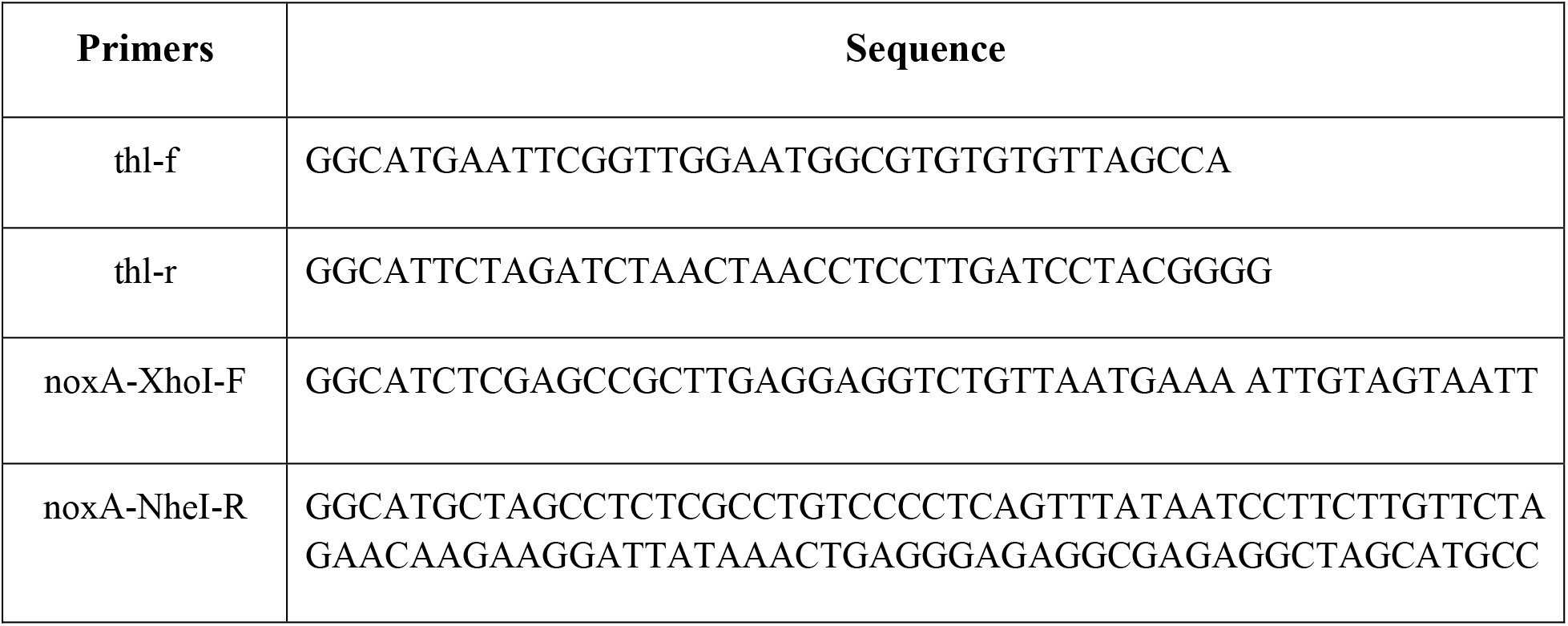
Primers used in PCR to construct the plasmid

The resulting plasmid, pMTLTN, was transformed into competent *E. coli* CA434 cells by electroporation using a Gene Pulser Xcell Electroporation System (Bio-Rad Laboratories Ltd, Mississauga, ON, Canada). The pre-set protocol for *E. coli* was used with a 2 mm cuvette. The plasmid was transferred into *C. sporogenes* by conjugation, following the procedure described by Theys et al. (Theys *et al*., 2006). Briefly, transformed donor *E. coli* was inoculated in LB broth supplemented with erythromycin. The culture was incubated overnight at 37° C in an orbital shaker at 225 rpm. The target *C. sporogenes* strain was inoculated in TYG media and incubated overnight at 37° C under anaerobic conditions. On the second day, 1 ml of the overnight *E. coli* donor culture was pelleted by centrifugation at 8000 rpm for 1 minute. The supernatant was discarded, and the cells were washed by re-suspending in 0.5 ml sterile PBS buffer. The centrifugation step was repeated, and the supernatant was again discarded. The donor *E. coli* pellet was re-suspended in 200 μl of the overnight *C. sporogenes* culture to produce a conjugation mixture. The entire conjugation mixture was pipetted onto a single non-selective TYG agar plate in discrete spots. The plate was incubated for 8 hours at 37° C in anaerobic conditions to allow conjugal transfer of the plasmid from the *E. coli* donor to the *C. sporogenes* recipient. One ml of anaerobic sterile PBS was pipetted onto the conjugation plate. Using a sterile spreader, the layer of cells was scraped off the plate and was re-suspended in PBS. Using a pipette, the cell-PBS slurry was transferred to a fresh micro-tube. Then 100 μl of slurry and its 10-fold dilutions were spread onto fresh plates containing TYG agar plate supplemented with 250 µg/ml D-cycloserine, to select against the *E. coli* conjugal donor, and 10 µg/ml erythromycin, to select for the plasmid. After incubation at 37° C for 24 hours, colonies were large enough to pick.

### 2.2 Bacterial Growth

Anaerobic growth of *Clostridium* strains occurred in a Thermo Forma Anaerobic System, model 1025, under an atmosphere of 5% carbon dioxide (CO_2_), 10% hydrogen (H_2_), and 85% nitrogen (N_2_), at 37° C, in TYG medium (3% trypticase, 2% yeast extract, and 0.1% glucose). Microaerobic growth occurred in a Plas-Labs Controlled Atmosphere Chamber (CAC) in a 5% CO_2_, 10% O_2_, and 85% N_2_ atmosphere, at 37° C. *E. coli* was grown in LB broth (1% trypticase, 0.5% yeast extract and 1% NaCl) and on LB agar (1.5% agar) at 37° C. For transformed *E. coli* strains, growth culture was supplemented with 500 μg ml^-1^ erythromycin. For *Clostridium sporogenes*, antibiotic concentrations were 5 μg ml^-1^ (erythromycin) and 100 μg ml^-1^ (D-cyclocerine). *E. coli* and *C. sporogenes* were stored at -80° C in glass cryovials containing 15% v/v and 10% v/v glycerol, respectively.

### 2.3 Real-Time Quantitative Reverse Transcription (qRT)-PCR Analysis

The native and PTN strains were cultured on TYGA plates (3% trypticase, 2% yeast extract, 0.1% glucose, and 1.5% bacterial agar). After 16 h, one colony of each strain was picked and used to inoculate 10 ml of TYG media. These cultures were harvested in exponential phase at OD_600nm_ 0.9. The cells were washed twice with TSE buffer (50mM Tris-HCl, 25 mM EDTA, 6.7% sucrose, pH of 8). Then 300 µl of TE buffer (50mM Tris-HCL, 20 mM EDTA, pH of 8) and 47 µl of freshly prepared lysozyme buffer (100 mg/ml lysozyme in RNase-free water) was added to the cells. The cells were resuspended by vortexing and the tubes were kept on ice for 10 minutes. Then, 500 µl of 10% SDS solution was added, and the solution was vortexed. The cells were incubated for 2 h at 37° C. All steps of the process were performed inside an anaerobic chamber. The remaining steps for binding, washing, and elution of the RNA from bacterial cell samples were carried out according to the PureLink® RNA Mini Kit instructions. The quality and yield of RNA were analyzed by the Take-3 Plate and Cytation 5 plate reader (BioTek). For the elimination of genomic DNA contamination, DNase I Amplification Grade was used (Invitrogen). The SuperScript™ First-Strand Synthesis System for RT-PCR was applied to synthesize the first strand of cDNA from purified RNA, according to the manufacturer’s instructions. Primer Quest Tool was used for primer and probe design (Integrated DNA Technologies Inc., Coralville, Iowa, USA). The analyzed genes, primers, and probes are listed in Table 2. qRT-PCR assays were performed with the TaqMan PCR Master Mix (Integrated DNA Technologies Inc.) on an Applied Biosystem 7500 Fast Real-Time PCR instrument according to the manufacturer’s instructions.

**Table 2-.**
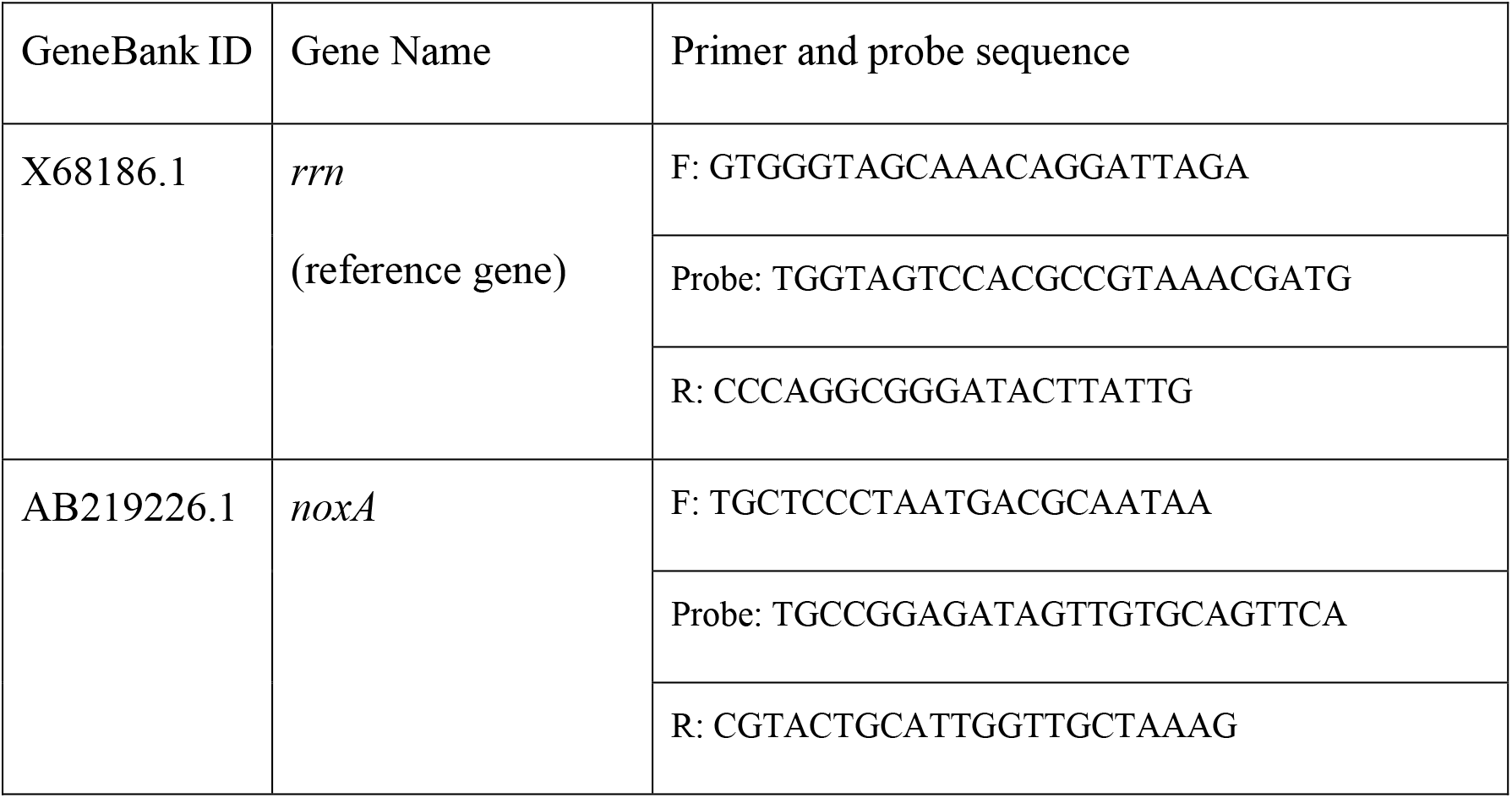
Genes and primers for real-time qRT-PCR

**Table 3-.**
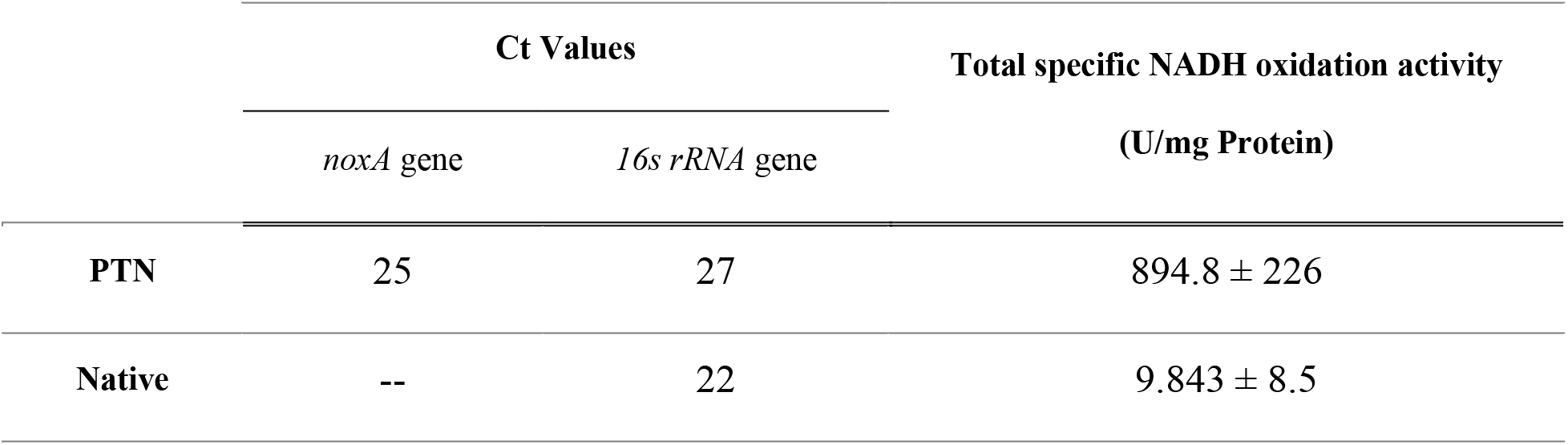
Gene expression analysis and NADH oxidation activity results

### 2.4 Enzyme Activity

#### 2.4.1 Harvesting and Lysis of Bacterial Cells

The native and PTN strains were cultured on TYGA plates. After 16 h, one colony of each strain was picked and inoculated in TYG media. The bacteria were harvested in the late log phase of growth, centrifuged at 10000 g for 15 min, and the pellet washed twice with 50 mM potassium phosphate buffer (pH 7.8) containing 0.1 mM EDTA. The final cell pellet was resuspended in 500 µl of 25 mM Tris, pH 7.4, containing 0.01 mM EDTA, 0.2% Triton X-100, 10 mg/ml lysozyme, and 1x protease inhibitor, and then incubated at 37° C for 30 minutes. The suspension was subjected to ultrasound treatment (Pulse 150 Ultrasonic Homogenizer, 10 cycles of 45 s ON and 1.5 min OFF at power rate of 40%) on ice. The homogenized suspension was centrifuged at 15000 rpm for 20 minutes at 4° C. The supernatant was used for the enzymatic assay, as described below. Total soluble protein was quantified using the Pierce™BCA Protein Assay Kit.

#### 2.4.2 Enzyme Activity

The activity of water-forming NADH oxidase was assayed spectrophotometrically in 1 ml of air-saturated 50 mM sodium phosphate buffer (pH 7.0) containing 0.15 mM NAD(P)H at 37° C, following the procedure described in (Kawasaki *et al*., 2005). The reaction was initiated by the addition of supernatant as described above. The decrease in absorbance at 340 nm was monitored with a spectrophotometer (BioTek). One unit of activity was defined as the amount of enzyme that catalyzes the oxidation of 1 µMol NAD(P)H per min.

### 2.5 Growth and Viability Assay

#### 2.5.1 Experimental Setup and Growth Condition

The native and PTN strains were streaked on TYGA plates which were incubated at 37° C in anaerobic conditions. After 24 h, one colony from each plate was picked to inoculate 10 ml of TYG culture media for overnight growth. The next day, subcultures of each of the two overnight cultures were prepared in TYG culture media. At mid-log phase, the cultures were diluted into either anaerobic media or oxygen-preconditioned media, as indicated below. Then 1.5 ml of each culture was aliquoted in a 24-well plate, in triplicate. These plates were prepared inside an anaerobic chamber (to avoid exposing the bacteria to atmospheric oxygen). To prepare preconditioned media, a bottle of TYG media covered by a cotton plug was put on a shaker in a Controlled Atmosphere Chamber (CAC) set to 10% environmental oxygen. The 24-well plate was transferred into the CAC using an anaerobic jar to avoid any atmospheric oxygen exposure. The cultures were incubated at 37° C for 48 h in a Cytation 5 plate reader (BioTek) inside the CAC. The plate reader was programmed to shake the plate at 180 cpm. During growth, 100 µl samples of each culture were collected at each time point to analyze viability and sporulation.

#### 2.5.2 Enumeration of Spores and Vegetative Cells

Duplicate samples were taken from the multi-well plate at each assay time-point. One sample from each pair was heat-treated at 70° C for 10 min to kill all vegetative cells. 10-fold serial dilutions of both heat-treated and non-heat-treated samples were prepared in sterile oxygen-free PBST (1X Phosphate-Buffered Saline, 0.1% Tween® 20 Detergent) in the anaerobic chamber. Then 100 µl of each dilution was placed on a TYG agar plate by spread plating. The plates were incubated anaerobically for 3 days at 37° C. Colonies were manually counted after 72 h of incubation. Following (Wang et al., 2017), total viable cell counts (CFU/ml) were determined from the samples that were not heat-treated; spore counts (CFU/mL) were determined from the heat-treated samples. The vegetative cell count was taken as the difference between the two.

### 2.6 Oxygen Concentrations in Liquid and Gas Phase

The dissolved oxygen concentration in TYG broth was measured using a Compact Extech DO210 Dissolved Oxygen Meter with a resolution of 0.1 ppm. The oxygen concentration in the gas phase of the environmental incubators was continuously measured using a LuminOx oxygen gas sensor with a resolution of 0.01%. The dissolved oxygen values were measured after equilibrium was reached under similar conditions used for the growth in multi-well plates.

## 3 Results

### 3.1 *noxA* Gene Expression and Enzyme Activity Assay

Transcriptional expression of the *noxA* gene and 16s rRNA (as the housekeeping control) were assayed by qPCR in both the PTN and native strains (Table 1), indicating that there was expression of the *noxA* gene in the PTN strain while there was no evidence of *noxA* expression in the native strain. To assess the activity of the expressed NoxA enzyme, the oxidation capacity of the cell lysate was measured (Table 1), revealing a significant difference between the activity from the PTN and native strains.

### 3.2 Growth of PTN and Native Strains in 10% Environmental O_2_: inoculation into Aerobic Media

Growth of the PTN and native strains were observed for two inoculum sizes, both inoculated from mid-log phase anaerobic cultures into TYG culture media that had been preconditioned in 10% environmental oxygen as described in Methods. The media was confirmed to contain 2.5 ± 0.1 mg/L dissolved oxygen (DO). As shown in Figure 1, the PTN strain had a growth advantage for both inoculum sizes, although the growth advantage is reduced for the smaller inoculum (panel B). A viability assay for PTN growth from the larger inoculum (panel A) showed 10^8^ CFU/ml in the stationary phase. Further investigation confirmed that the native strain formed a biofilm in these conditions [Supplementary Figure 1].

**Figure 1-.**
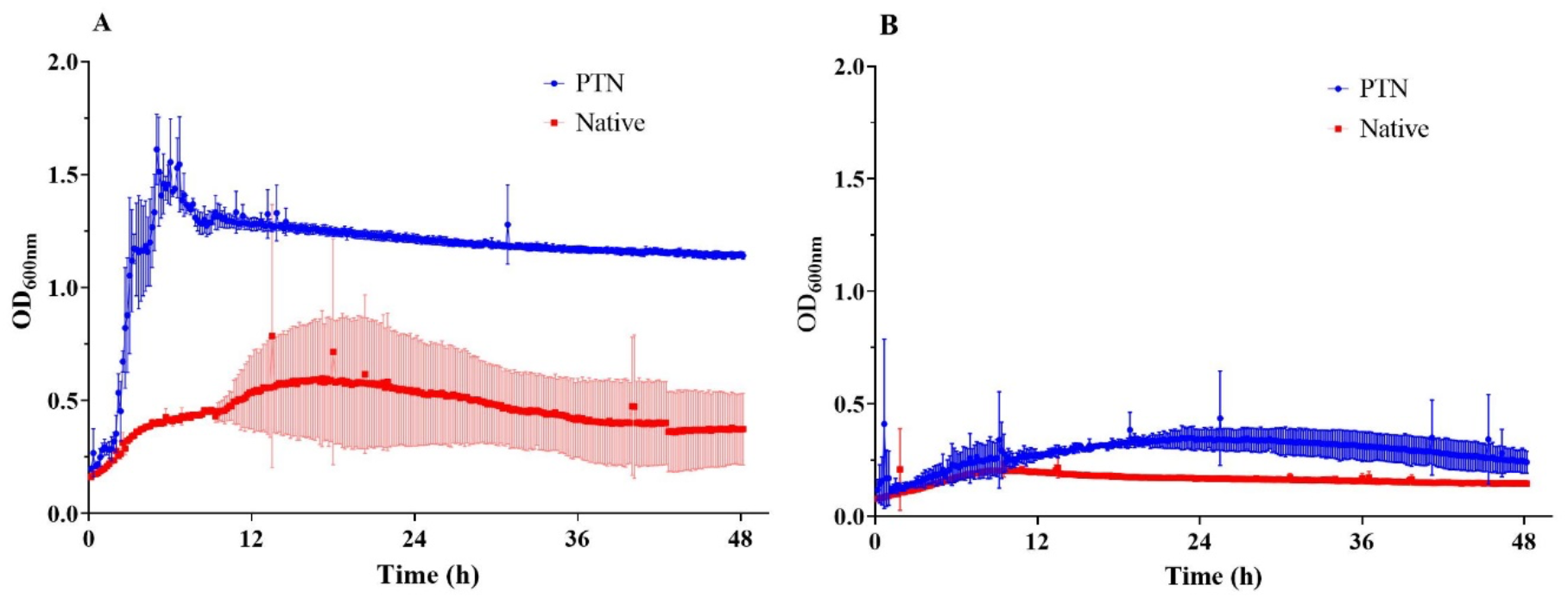
Growth of the PTN and native strains in preconditioned TYG medium, containing 2.5 mg/L dissolved oxygen, in 10% environmental oxygen. Inoculum density OD_600nm_ 0.2 (A), and OD_600nm_ 0.1 (B). The *noxA*-expressing PTN strain shows improved growth in both cases, but the smaller inoculum (panel B) provides a much smaller growth advantage. Error bars indicate one standard deviation from triplicate cultures.

### 3.3 Growth of PTN and Native Strains in 10% Environmental O_2_: inoculation into Anaerobic Media

To investigate the transition in growth capacity as oxygen stress increases, the PTN and native strains were inoculated into anaerobic TYG media (not pre-exposed to oxygen) and then grown in 10% environmental oxygen. Here, the oxygen level in the culture media increases through time until saturation with the environment is reached. As shown in Figure 2A, the initial growth rate for the PTN and native strains is similar. After 8 h, the optical density (600 nm) of the native strain began to decline. In contrast, the PTN strain maintained a steady OD, suggesting a prolonged stationary phase. Spore counts (Figure 2A, representative images in panels B and C) show that the PTN strain maintained itself in a vegetative state throughout the experiment, while the native strain began sporulation after 8 h. After 48 h, almost all of the native cells were in spherical spore-form.

**Figure 2-.**
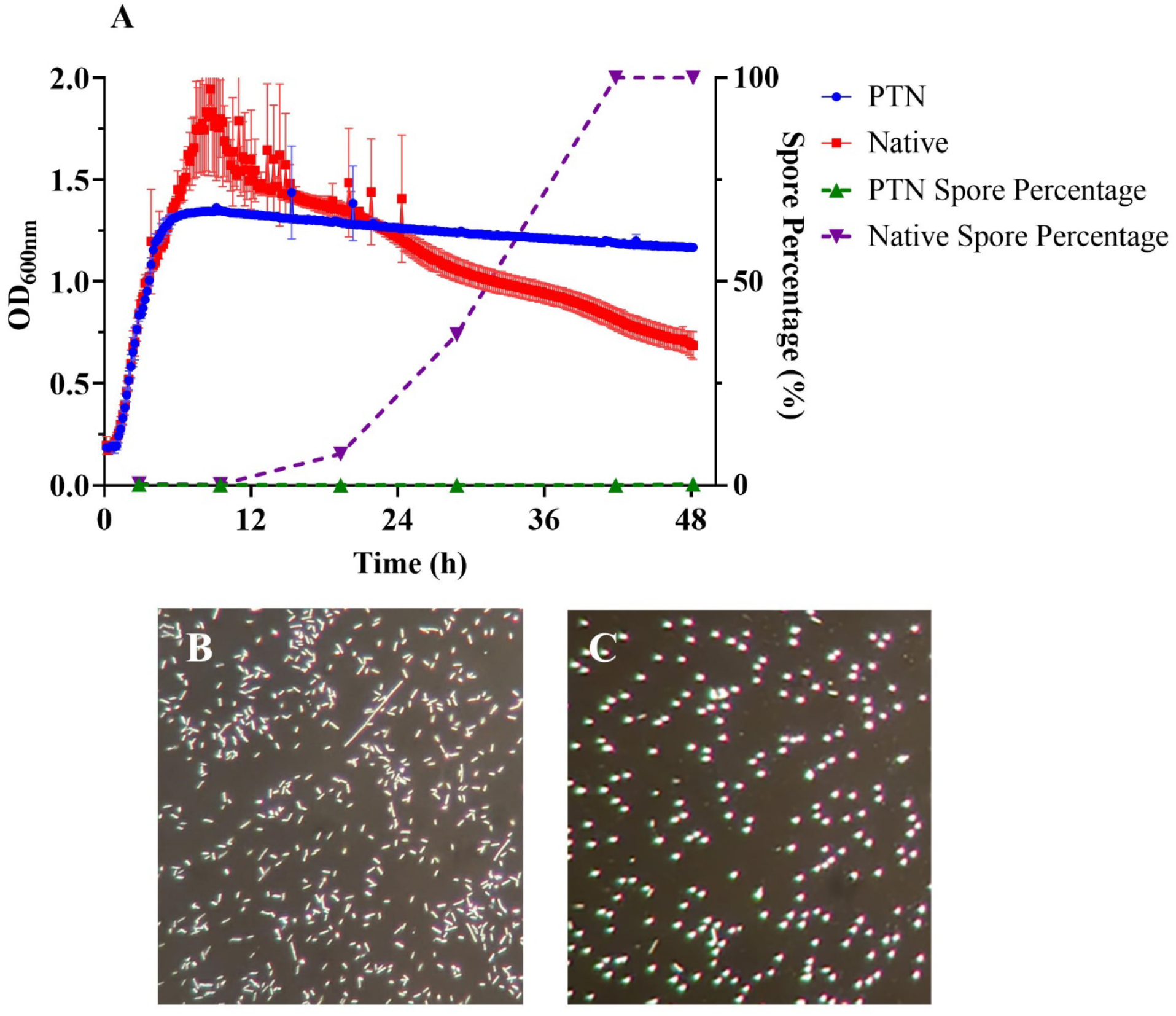
Growth of the PTN and native strains in 10% environmental oxygen, cultured in initially anaerobic media. (A) Growth curve and spore enumeration of the PTN and native strains. Initially, growth was similar. The native strain had a short-lived stationary phase before the OD began to decline. In contrast, the PTN strain had a prolonged stationary phase in which the number of vegetative cells was constant (∼10^8^ CFU/ml). After 8 h, the native strain began sporulation, with the majority of the population in spore form by 48 h. (B) Microscopic observation of the PTN cells after 48 h. The cells are in vegetative, rod-shape form; (1000x magnification). (C) Microscopic observation of the native cells after 48 h. The cells are mostly spherical spores; (1000x magnification). Enumeration of spores (panel A) was performed by heat-treatment to eliminate vegetative cells (details in Methods). Error bars indicate one standard deviation from triplicate cultures.

### 3.4 Effect of Inoculum Size on Growth

To further investigate the effect of inoculum size exhibited in Figure 1, we investigated growth from a range of inoculum sizes inoculated in anaerobic TYG media and cultured in a 10% oxygen environment (Figure 3). As in Figure 2, this protocol allows for a period of growth before the culture media became saturated with oxygen. Figure 3A shows that, as in Figure 2, at a high inoculum size, both strains grew initially, but as oxygen levels in the media rose, the native strain was unable to maintain vegetative viability, while the PTN strain remained viable through a stationary phase. From a smaller inoculum (panel B), the native strain showed no growth, while growth of PTN occurred after a prolonged lag phase. When the inoculum was even smaller, (panel C) neither strain grew.

**Figure 3-.**
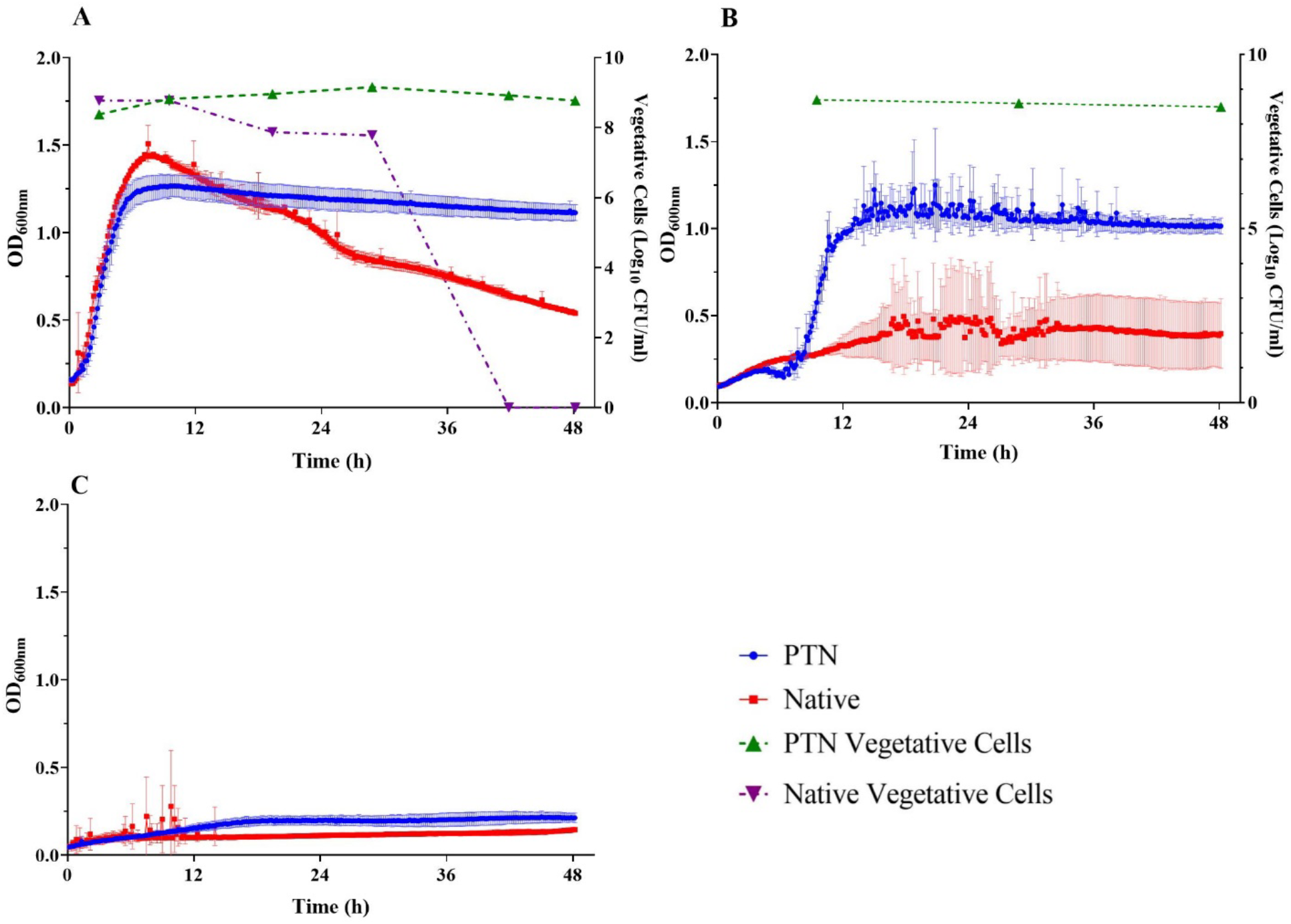
Growth of PTN and native strains in 10% environmental oxygen when inoculated into anaerobic media. All tubes were inoculated from a culture at 1.6 OD_600_. Growth of the PTN and native strains from inoculum at 12% (v/v) (A), 6% (v/v) (B), and 3% (v/v) (C). Error bars indicate one standard deviation from triplicate cultures. The largest inoculum (panel A) led to significant growth of both strains, but vegetative viability of the native strain began to decline after 8 h, while PTN viability was maintained in stationary phase. At the middle inoculum (panel B), PTN growth occurred after a significant lag, while the native strain showed minimal growth. At the lowest inoculum (panel C), neither strain showed growth. (Viability tests were not performed for cultures that did not show growth.)

## 4 Discussion

In the present study, we introduced the *noxA* gene, which codes for a water-forming NADH oxidase, into *C. sporogenes* to develop an aerotolerant strain. This engineered PTN strain was observed to grow to OD of about 1.2 and retain vegetative viability for at least 48 h at 10% environmental oxygen, which, in saturation, is equivalent to 2.5 mg/L dissolved oxygen in the culture media. In contrast, in the same conditions, the parental native strain grew only to OD 0.4. Moreover, the native strain produced a biofilm over the course of the experiment: a response that may provide a different measure of oxygen tolerance. Biofilm formation has been observed in the R20291 *C. difficile* strain as a protective mechanism against oxygen stress (Dawson et al., 2012), and likewise has been observed in *C. perfringens* in response to H_2_O_2_ stress (Varga, Therit and Melville, 2008). In addition, there may be intrinsic mechanisms for oxygen tolerance at play, aiding the native culture in these conditions, as has been confirmed for *C. botulinum* (Camerini *et al*., 2019). Studies have shown that *Clostridia* can cope with oxygen better than might be expected for obligate anaerobes (Lu and Imlay, 2021; Morvan *et al*., 2021). However, *C. sporogenes* has been observed to be more sensitive to oxygen than other *Clostridia* species (Kawasaki *et al*., 1998).

When the culture media was equilibrated in an anaerobic environment prior to inoculation transfer to a 10% environmental oxygen chamber (Figure 2A), both strains initially grew well. However, over time, as oxygen diffused into the medium, the OD of the native strain started to decrease, which we further found to correlate with sporulation. Sporulation in response to oxygen stress has been observed in a range of *Clostridium* species (Al-Hinai, Jones and Papoutsakis, 2015; Diallo, Kengen and López-Contreras, 2021).

As observed in Figure 1A, the degree of aerotolerance of the engineered PTN strain is dependent on inoculum size, with the smaller inoculum less able to tolerate oxygen stress. We hypothesize that this effect is a result of insufficient NoxA production from small populations. We explored this population-size effect further, as shown in Figure 3. At the highest inoculum size both strains grew, but only the PTN strain maintained vegetative viability. From a smaller inoculum, PTN growth was delayed, and each strain exhibited a threshold inoculum size below which growth was not observed. This population-size effect suggests that secretion of protective factors (including NoxA in the case of PTN) was responsible for defense from oxygen stress.

*Clostridia* spp. are promising agents for cancer therapy: they have the selective capability to target solid tumors and can be used to deliver therapeutic factors (Mowday *et al*., 2016). However, their use in cancer therapy is limited by their inability to penetrate the outer oxygenated parts of a tumor, allowing tumor regrowth (Mengesha *et al*., 2007; Mowday *et al*., 2016). The aerotolerant PTN strain presented here could be a promising vehicle for clostridia-mediated tumor therapy. The oxygen tolerance conferred by NoxA maintains *C. sporogenes* in vegetative form in 2.5 mg/L dissolved oxygen in media (pO_2_ ∼ 7 %, details in Supplementary Material). This level of dissolved oxygen is consistent with the oxygenated parts of a tumor (pO_2_ ∼ 7-8 %) (Carreau *et al*., 2011). Oxygen levels through solid tumors and surrounding healthy tissue vary widely across tumor and tissue types (Feldmann, Molls and Vaupel, 1999; Carreau *et al*., 2011); promoter tuning could be used to tailor strains to target regions with specific oxygenation profiles. As demonstrated above, the NoxA-derived aerotolerance is not conferred at small population sizes. Consequently, individual spores of this engineered strain would be unable to germinate in oxygenated tissue, and so this strain would retain the tumor-targeting features of *C. sporogenes*, enabling it to selectively colonize and germinate in the necrotic and hypoxic region of solid tumors (pO_2_ ≤ 1%-1.3 %). Further refinement of this strain could result in appropriate regulation of *noxA* expression to avoid toxicity or damage to healthy tissue, e.g. by introducing an environment-sensitive promoter.

Oxygen tolerance in *Clostridia* may also be beneficial in biomanufacturing, where these strains are used in a number of processes, such as ABE fermentation (focused on *C. acetobutylicum*) and other bioproduction systems for chemicals and fuels (Moon *et al*., 2016; Zeng, 2019; Zhao *et al*., 2020). Focusing specifically on *C. sporogenes*, this strain has been used for non-acetone forming butanol production from rice straw (Gottumukkala *et al*., 2013, 2015) and has recently been identified as a producer of the potent antioxidant 3-indolepropionic acid (IPA) (Du *et al*., 2021), which has neuroprotective properties (Zhang and Davies, 2016). The use of *Clostridium* has been limited in these contexts due to the requirement of an oxygen-free environment. Development and maintenance of a strict anaerobic environment demands expensive and complex equipment and operations (Papoutsakis, 2008). For example, in syngas fermentation by acetogens (including *Clostridium*), oxygen is a common contaminant, inhibiting acetogen growth and product formation (Daniell, Köpke and Simpson, 2012). Attempts have been made to overcome oxygen limitation through the development of *Clostridium* co-culture systems to control the extracellular environment (Du et al., 2020; Ebrahimi et al., 2020; Cui, Yang and Zhou, 2021). A tunable aerotolerant strain of *Clostridia*, as demonstrated here, could offer a new angle for meeting these challenges.

## 4 Author Contributions

Project design: BI, BZ, SS, MGA, JP. Experiments: SS, BZ, Manuscript draft: BI, SS, MGA Manuscript editing: BI, SS, MGA

