## Supplement for "Heterologous Expression of NoxA Confers Aerotolerance in *Clostridium sporogenes*"

### Supplementary Material

#### 1 Supplementary Figures

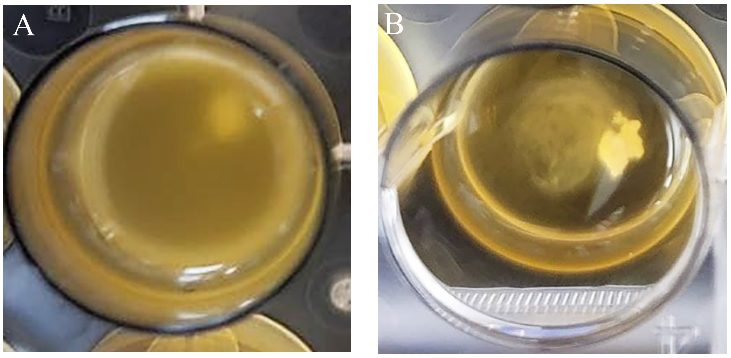


**Supplementary Figure 1.** Liquid culture of the PTN and native strains in preconditioned TYG medium containing 2.5 mg/L dissolved oxygen, in 10% environmental oxygen after 48 h incubation. Inoculum density is $OD_{600nm}$ 0.2. (A) Liquid culture of PTN showing growth (turbidity). (B) Biofilm formation by the native strain.

#### 2 Supplementary Data

Solubility as a function of temperature and Henry's constant (H) at 25º C for gases in water can be determined from: (Green & Southard, 2018, p.2-89)

$$\ln x = A + \frac{B}{T} + C \ln T + DT$$

where, T is temperature (K), x is mole fraction of the solute dissolved in water when the solute partial pressure is 1 atm, and the constants of A, B, C, and D, in appropriate units, for oxygen as a solute in water are:

$$A = -171.2542$$

$$B = 8391.24$$

$$C = 23.24323$$

$$D = 0$$

In our experiments, $T = 310$ K.

The solubility of oxygen in water at 37º C can be calculated as follows:

$$\ln x = (-171.2542) + \left(\frac{8391.24}{310}\right) + 23.24323 \times \ln(310) = -10.85$$

which gives:

$$x = 1.95 \times 10^{-5}$$

We then have Henry's constant at 37° C as

$$H = \frac{1 \text{ atm}}{x} = 51546.4 \text{ atm}$$

To convert the value of H from atm to $\frac{\text{atm}}{\frac{\text{mol}}{\text{m}^3}}$, we divide by the molar density of water, which is $55342 \ \frac{\text{mol}}{\text{m}^3}$, giving

$$H = 0.931 \ \frac{\text{atm}}{\text{mol/m}^3}$$

This is the Henry constant for oxygen in water at 37° C.

In our experiment, we measured dissolved oxygen in media as $c_{O_2} = 2.5 \text{ mg/L}$.

Based on Henry's law, the partial pressure of oxygen in water, $p_{O_2}$, is proportional to the molar concentration of oxygen in water:

$$p_{O_2} = Hc_{O_2}$$

The molecular weight of oxygen is 32 g. Thus, the concentration of oxygen in our media is:

$$2.5 \frac{\text{mg}}{\text{L}} \times \frac{1 \text{ mol}}{32000 \text{ mg}} \times \frac{1 \text{ L}}{0.001 \text{ m}^3} = 7.81 \times 10^{-2} \ \frac{\text{mol}}{\text{m}^3}$$

The partial pressure of oxygen in the media thus:

$$p_{O2} = 0.931 \ \frac{\text{atm}}{\frac{\text{mol}}{\text{m}^3}} \times 7.81 \times 10^{-2} \ \frac{\text{mol}}{\text{m}^3} = 0.0727 \text{ atm}$$

In SI, the pressure unit is the Pascal (Pa). Other units include the atmosphere (1 atm = 101.325 kPa), and the bar (1 bar = 100 kPa). Two other units, mostly used in medicine, are the millimetre of mercury (1 mmHg = 133.322 Pa) and the percentage of oxygen (1% = 1.013 kPa), (Carreau et al., 2011).

The 2.5 $mg/L$ oxygen dissolved in water is equivalent to partial pressure ($p_{O2}$) of 7.27%.

The chemical properties and salinity of media differ from water, which affects the value of Henry's constant. However, we can conclude that the approximate value of partial pressure of oxygen ($p_{O2}$) in the media is ~ 7%.
